# Cortical ensemble activity discriminates auditory attentional states

**DOI:** 10.1101/622332

**Authors:** Pan-tong Yao, Jia Shen, Liang Chen, Shaoyu Ge, Qiaojie Xiong

## Abstract

Selective attention modulates sensory cortical activity. It remains unclear how auditory cortical activity represents stimuli that differ behaviorally. We designed a cross-modality task in which mice made decisions to obtain rewards based on attended visual or auditory stimuli. We recorded auditory cortical activity in behaving mice attending to, ignoring, or passively hearing auditory stimuli. Engaging in the task bidirectionally modulates neuronal responses to the auditory stimuli in both the attended and ignored conditions compared to passive hearing. Neuronal ensemble activity in response to stimuli under attended, ignored and passive conditions are readily distinguishable. Furthermore, ensemble activity under attended and ignored conditions are in closer states compared to passive condition, and they share a component of attentional modulation which drives them to the same direction in the population activity space. Our findings suggest that task engagement changes sensory cortical representations across modalities in the same directions, and cross-modality attention may differentially modulates attended and ignored modalities.

## Introduction

Sensory perception is highly modulated by attention (Petersen and Posner 2012). At different modes and levels of engagement in behavioral tasks, attention may modulate sensory cortical processing, including spontaneous activity (Arieli, Sterkin et al. 1996, Kastner, Pinsk et al. 1999, Buran, von Trapp et al. 2014, Rodgers and DeWeese 2014), stimulus-evoked activity (Hubel, Henson et al. 1959, Fritz, Shamma et al. 2003, Otazu, Tai et al. 2009, Carcea, Insanally et al. 2017), and population dynamics (Womelsdorf, Fries et al. 2006, Cohen and Maunsell 2009, Briggs, Mangun et al. 2013). Sound representations in the auditory cortex change in response to the activation of neuromodulatory systems that regulate attention (Kilgard and Merzenich 1998, Bao, Chan et al. 2001, Marlin, Mitre et al. 2015, Martins and Froemke 2015). Depending on behavioral contexts, the same stimulus can be a target that requires attention or a distractor that should be ignored. Here, we examine whether auditory cortical neurons, at both single-cell and population levels, respond to stimuli differently when they are targets versus when they are distractors in a cross-modality attention task, and how ensemble neuronal activities differ under these attentional conditions.

## Results

### Behavioral and recording paradigms for mice attending to or ignoring the same auditory stimuli

To determine whether auditory cortical neurons respond differently to the same stimuli under attended or ignored conditions, we designed a cross-modality attention task (**Fig. 1A–C**). We first trained mice to perform a two-alternative forced choice (**2AFC**) sensory discrimination task. In brief, a freely moving mouse was placed in a dark, sound-proof chamber. Each trial was self-initiated by the mouse poking its nose into the center port to trigger a sound and/or light stimulus. In the sound block, a stream of pure tones with different frequencies was presented as the cue. The mouse learned to associate the frequency of pure tones (high versus low) with an action (going to the left or right port) for a water reward (**Fig. 1A**). In the light block, LED lights on top of either left or right port were turned on as a cue, and a stream of pure tones with different frequencies was simultaneously presented as a distractor (i.e., their frequencies were not associated with the reward port). The mouse learned to go to the lit port for the water reward and ignore the auditory distractor (**Fig. 1B**). Well-trained mice performed with average accuracies of 87.0 ± 1.6% in sound blocks and 91.2 ± 1.5% in light blocks (**Supplementary Fig. 1A**). In light blocks, because tone streams with either low or high frequencies were randomly assigned to each trial, there were concordant trials in which the tone frequency and the reward port indicated by the light had the same association as the sound blocks; discordant trials were those in which the tone frequency and the reward port had the opposite association as the sound blocks. Mice performed with accuracies of 99.4 ± 0.3% in concordant trials and 85.6 ± 2.2% in discordant trials (**Supplementary Fig. 1B**). Together the behavioral results showed that mice learned to attend to the auditory targets in sound blocks and ignore the auditory distractors in light blocks (**Fig. 1D**).

**Figure 1.**
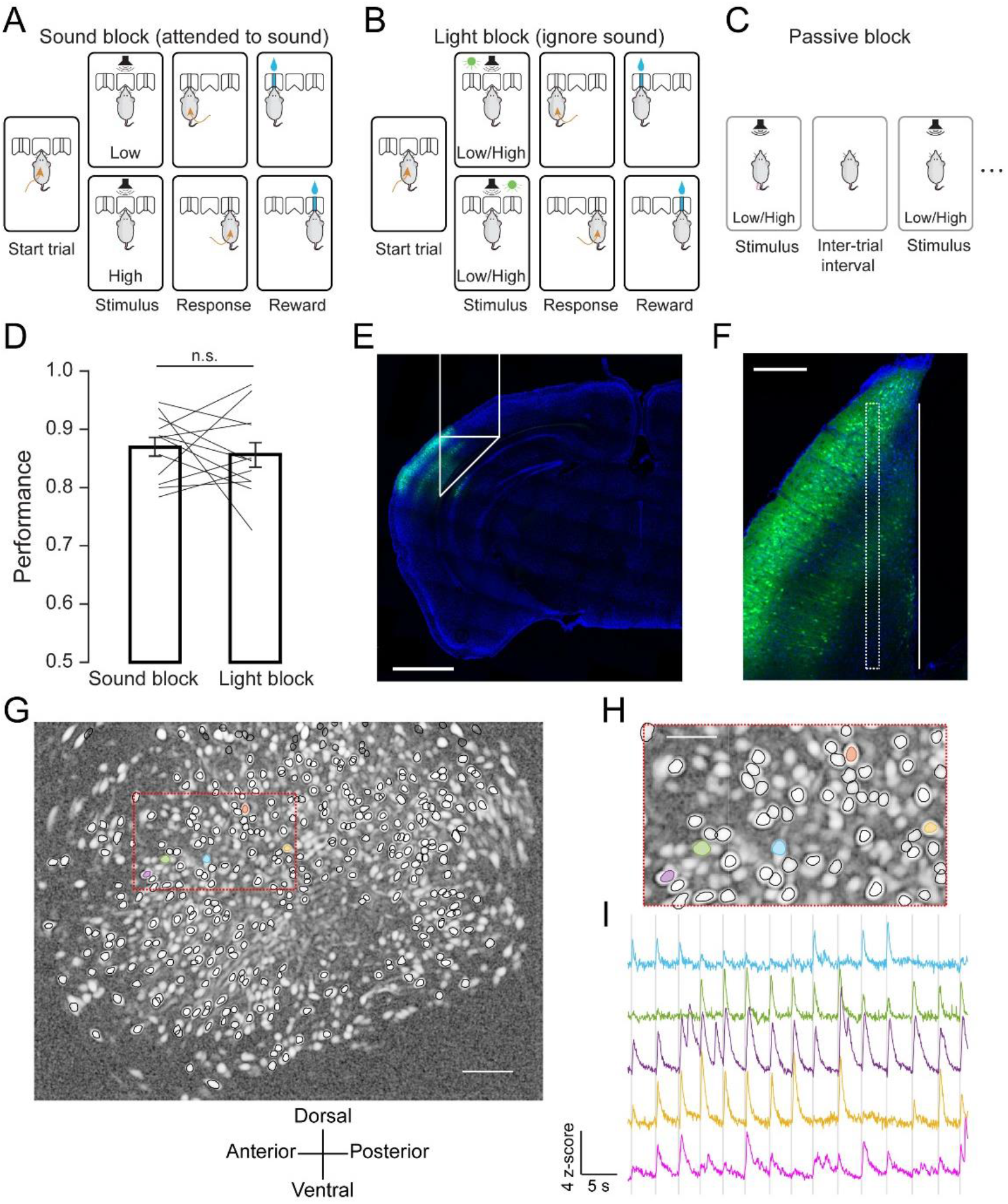
Behavioral paradigm and in vivo calcium imaging of auditory cortical neurons. **A.** Illustration of the behavioral task in the sound block, in which auditory stimuli were associated with rewards. **B.** Illustration of the behavioral task in the light block, in which visual (but not auditory) stimuli were associated with rewards. **C.** Passive block in which mice passively listened to an auditory stimulus. **D.** Task performance of mice (n = 5 mice, 12 sessions) in sound blocks (0.87 ± 0.02) and light blocks (only discordant trials are included: 0.86 ± 0.02); *P* = 0.64, paired-sample *t* test; data are presented as the mean ± SEM. **E.** AAV9-CaMKII-GCaMP6f expression and prism probe position in the auditory cortex. GCaMP6f: Green; DAPI: Blue. Scale bar: 1 mm. **F.** Coronal section from a representative mouse brain showing the prism probe tract with its imaged side facing the GCaMP6f-expressing cells. Solid line, prism probe tract; dashed line, focal plane. Scale bar: 200 μm. **G** & **H.** The contours of detected neurons superimposed on the image of a representative field of view. Scale bars: 100 μm (**G**), 50 μm (**H**). **I.** Fluorescence traces of the example region of interest (colored in **G** & **H**). Gray shading, sound presentation period.

We next recorded neuronal activity of the primary auditory cortex from well-trained mice using in vivo Ca^2+^ imaging. We expressed GCaMP6f, an ultrasensitive Ca^2+^ sensor protein (Chen, Wardill et al. 2013), in the primary auditory cortex by the stereotaxic injection of adeno-associated virus (AAV) and then implanted a prism lens above the injection site for Ca^2+^ imaging using miniaturized fluorescence microscopy, as described previously (Kirschen, Shen et al. 2017) (**Fig. 1E&F**). GCaMP6f is controlled by the CaMKII promotor; therefore, we monitored excitatory neurons in the primary auditory cortex. Four weeks after viral infection, we imaged Ca^2+^ activity from these mice when tasks were performed in sound blocks and light blocks, or when the mice passively listened to auditory stimuli (passive blocks, **Fig. 1C**). The Ca^2+^ signals from one recording session are shown in **Figure 1G–I**. We detected 3,379 neurons from 5 mice in 12 recording sessions. Among the detected neurons, 249 showed robust responses to the tone-cloud stimuli in the passive block (bootstrap, *P* < 0.01). These neurons are referred to as stimulus-responsive neurons below.

### Populational activity auditory cortical neurons differentiates different attentional states

To study the attentional modulation of neuronal activity from individual stimulus-responsive neurons, we compared the peak intensity of the average calcium trace of each neuron in a 0–500-ms time window from the onset of sound in three contexts: in the sound block when mice attended to the auditory stimuli, in the light block when mice ignored the auditory stimuli, and in a passive session when mice passively heard the stimuli (**Fig. 2A&B**). To avoid day-to-day variations, we performed comparisons of the three contexts from sessions recorded on the same day. From stimulus-responsive neurons, we observed both enhancement and suppression of evoked responses under attended condition compared to passive condition. The same bidirectional modulation of the stimulus-evoked responses was also observed under ignored condition (**Fig. 2A&B**). The similarities in the modulation index distribution of attended vs. passive and ignored vs. passive modalities suggest that engaging in the task modulates cortical neuronal activity under both attended and unattended conditions.

**Figure 2.**
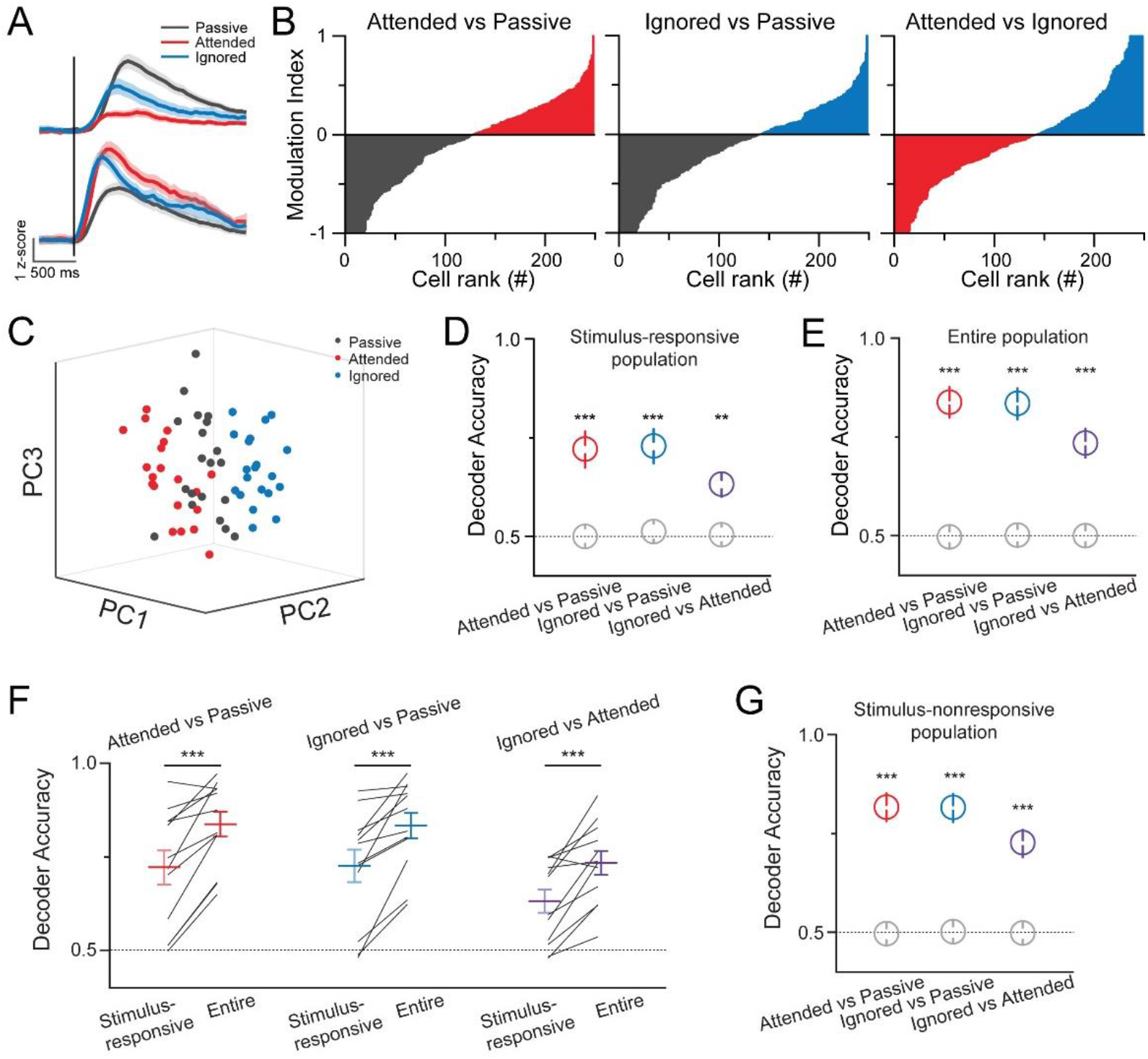
Activity of the auditory cortical neuron population distinguishes different attentional states. **A.** Example traces of evoked responses from two auditory cortical neurons to the auditory stimulus in the sound block (attended, red), light block (ignored, blue), and passive block (passive, grey). Black bar: onset of the auditory stimulus. **B.** The modulation index of individual neurons was compared between the three different attentional states. Modulation index: Calcium signal peak amplitudes (condition 1 – condition 2)/(condition 1 + condition 2). Cells were sorted based on the modulation index from −1 to 1 for each panel. (n = 249, grey: passive-preferring neurons, red: attended-preferring neurons, blue: ignored-preferring neurons.) **C.** A plot of population activity of stimulus-responsive neurons immediately after stimuli onset in different attentional states (20 trials for each state). Dots represent individual trials and are plotted in the same dimensionality-reduced space. **D** & **E.** Decoder performance of the stimulus-responsive population (**D**) or entire recorded population (**E**) in distinguishing two attentional states after stimuli onset: attended versus passive (red, *P* = 5.14 × 10^−4^, *P* = 6.90 × 10^−7^), ignored versus passive (blue, *P* = 4.33 × 10^−4^, *P* = 8.41 × 10^−7^), ignored versus attended (purple, *P* = 1.9 × 10^−3^, 1.50 × 10^−5^). Colored circles are from experimental data, and grey circles are from shuffled data. Statistical analyses were performed in a paired-sample *t* test between experimental and shuffled data (n = 12). All data represent the mean ± SEM. **F.** Comparisons of decoder performance between the stimulus-responsive population and the entire population after stimuli onset. (attended versus passive, red, *P* = 3.10 × 10^−4^; ignored versus passive, blue, *P* = 7.75 × 10^−4^; ignored versus attended, purple, *P* = 7.20 × 10^−4^; paired-sample *t* test, n = 12). **G.** Decoder performance of the stimulus-nonresponsive population in distinguishing two attentional states after the cue onset: attended versus passive (red, *P* = 4.68 × 10^−7^), ignored versus passive (blue, *P* = 5.08 × 10^−7^), ignored versus attended (purple, *P* = 1.43 × 10^−5^). Colored circles are from experimental data, and grey circles are from shuffled data. Statistical analyses were performed in a paired-sample *t* test between experimental data and shuffled data (n = 12). Data are presented as mean ± SEM.

The auditory cortical neuronal activity also displayed bidirectional differences in evoked responses between attended and ignored conditions (**Fig. 2B**). We asked whether cortical ensemble activity in response to targets could be distinguished from those in response to distractors, as well as under passive conditions. To quantify the differences in ensemble activity between attended, ignored, and passive conditions, we employed a support vector machine (SVM) as a decoder to analyze ensemble activity (See **Methods**). The decoder accuracy reflects how well the ensemble activity can distinguish different conditions (attended vs passive, ignored vs passive, attended vs ignored). The decoder accuracy is significantly above the chance level for all pairings after the onset of sound stimuli, which can be visualized in a dimensionality-reduced space (**Fig. 2C&D**). The results of engaged versus passive states (attended vs. passive, ignored vs. passive) indicated that the stimulus-responsive ensemble responds to the same auditory stimuli differently, with or without task engagement. Furthermore, the decoding results of the attended vs. ignored conditions showed that the stimulus-responsive ensemble differentially responds to the same auditory stimuli, depending on whether they are targets or distractors in the task context.

We further analyzed neuronal ensemble activity by applying the decoder to all recorded neurons. The decoder accuracies of the entire population were significantly higher than the chance level after the onset of sound stimuli (**Fig. 2E**), indicating that ensemble activity from the entire recorded neuronal population can distinguish attentional states. Interestingly, the decoder performance from the entire population was significantly higher than that from the stimulus-responsive population (**Fig. 2F**), suggesting that stimulus-nonresponsive neuronal activity also contributes to state separation. We thus performed the same analyses of the stimulus-nonresponsive population. Indeed, decoder accuracies were significantly higher than the chance level after the onset of sound stimuli (**Fig. 2G**). The fact that stimulus-nonresponsive neurons can distinguish the three attentional states suggests that these neurons are also modulated by task engagement and selective attention.

### Neuronal ensemble activity is in closer states between attended and ignored conditions

Decoder accuracies from the three neuronal populations in distinguishing attended (or ignored) from passive conditions were higher than that for attended vs. ignored (**Fig. 3A**), suggesting that the activity patterns are closer between the attended and ignored sessions than between the attended (or ignored) and passive sessions. We next determined whether selective attention during the performance of a task in the sound and light blocks modulated the ensemble activity in the same direction. Hypothetically, if attention modulates the cortical responses to targets and distractors in opposing ways, the modulation vectors from passive to attended states will have angles of 180° with the one from passive to ignored states (**Fig. 3B**, left panel), but if the attentional modulation of targets and distractors is in the same direction, the angle will be 0° (**Fig. 3B**, right panel), and if the attentional modulation of targets and distractors is independent, the angle will be 90° (**Fig. 3B**, middle panel). Our analysis showed that the angle between the modulation vectors of the stimulus responsive ensemble was 47.32 ± 5.48°, which is significantly different from that of the shuffled data (**Fig. 3C**, left panel, *P* = 4.35 × 10^−4^). The angles from the stimulus-nonresponsive ensemble and the entire ensemble were 56.78 ± 1.83° (**Fig. 3C**, middle panel, *P* = 5.67 × 10^−8^) and 54.78 ± 2.08° (**Fig. 3C** right panel, *P* = 1.19 × 10^−7^), respectively. These results indicate that the modulations of cortical ensemble activity under attended and ignored conditions share components that drive the population activity in the same direction.

**Figure 3.**
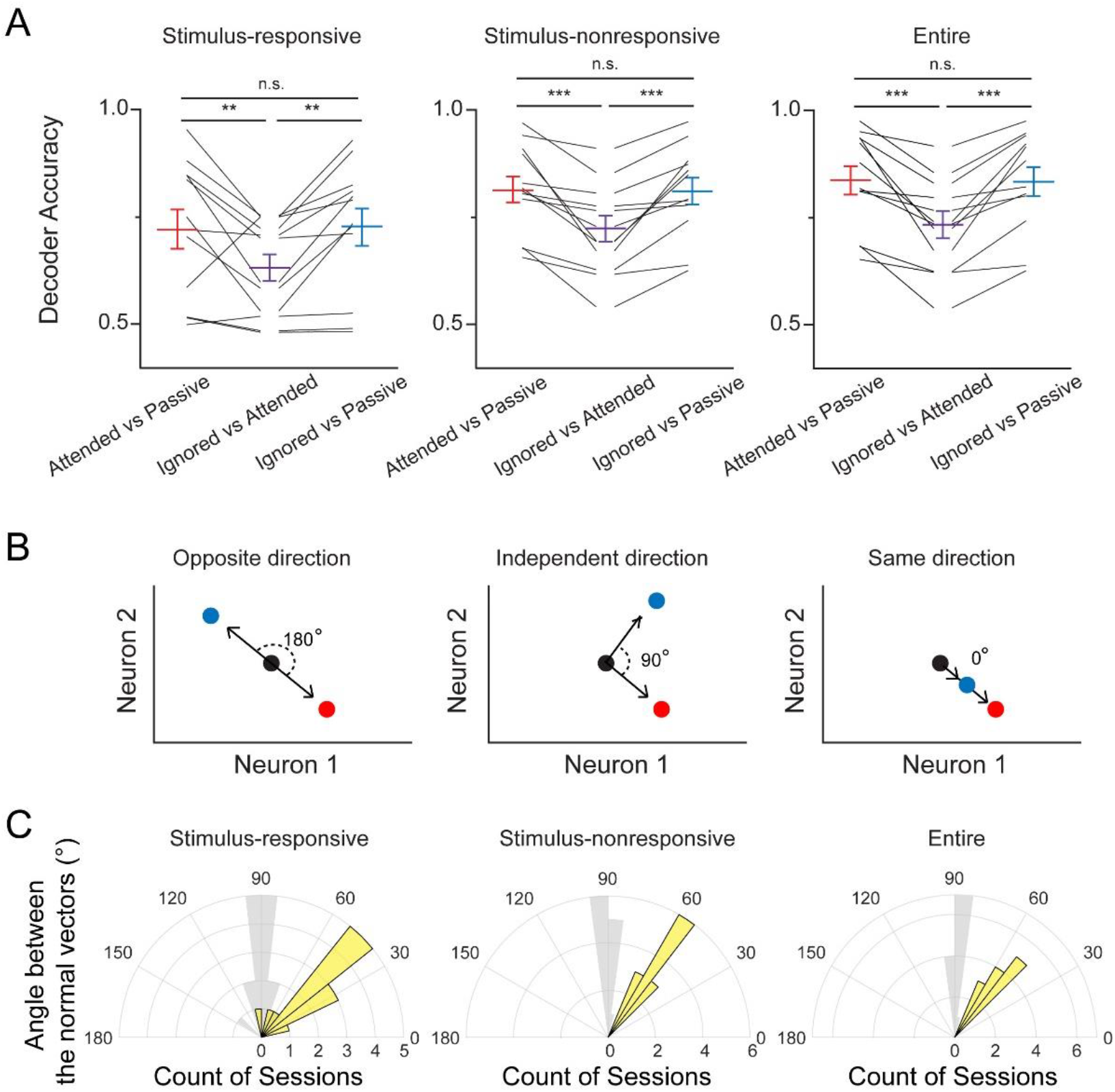
Attending and ignoring modulate population activity in the auditory cortex in the same direction. **A.** Comparisons of decoder performance between the three attentional states after stimuli onset. Stimulus-responsive population (left); stimulus-nonresponsive population (middle); entire population (right). Statistical analyses were performed in a one-way repeated measures analysis of variance followed by Tukey’s test. n.s., not significant. **B.** Hypothetical models of the directions of attentional modulation using example of two neurons. Black dots: passive states; blue dots: ignored states; red dots: attended states. **C.** Distribution of the angle between the normal vectors of the separate hyperplanes. Stimulus-responsive population (left); stimulus-nonresponsive population (middle); entire population (right). Yellow bars are from experimental data, and grey bars are from shuffled data.

## Discussion

Our study showed that the same auditory stimuli elicited different cortical activities when mice were performing the **2AFC** task compared to when they were passively listening to the stimuli (**Fig. 2A&B**). This finding suggests that engaging in the task induces attentional modulation of both attended and unattended sensory cortices. Multisensory spread of attention during modality-specific attention behavior has been reported in human studies (Mozolic, Joyner et al. 2008, Zimmer, Itthipanyanan et al. 2010), but the mechanisms responsible remain elusive. Both cholinergic innervation from the basal forebrain and noradrenergic innervation from the locus coeruleus to the neocortex are known to modulate cortical sensory representations in a behavior-dependent manner (Lin, Brown et al. 2015, Nelson and Mooney 2016, Kuchibhotla, Gill et al. 2017, Vazey, Moorman et al. 2018). Such neural inputs may be potential candidates for the circuitry mechanisms of the multisensory spread of attentional modulation.

Our results demonstrated that auditory cortical neurons respond to the same stimuli differently, depending on whether they are targets or distractors (**Fig. 2**). In addition to cross-modality modulation by engaging in behavioral tasks, there is sensory-selective modulation the attended and ignored modalities. Similarly, the circuitry mechanisms underlying such modulations remain unclear. Both cholinergic and noradrenergic signals, and the regulatory inputs from both the parietal and prefrontal cortices (Wimmer, Schmitt et al. 2015, Song, Kim et al. 2017), may play essential roles here, requiring further study.

Our analysis also showed that the ensemble activity of stimulus-nonresponsive neurons distinguished the attended, ignored, and passive states (**Fig. 2G**). Both the multisensory spread and modality-specific attentional modulation of these stimulus-nonresponsive neurons may change the local connections of stimulus-responsive neurons, which in turn may modulate the sensory processing that is important for relevant behaviors.

Our analysis suggests that cortical neuronal ensemble activity is in closer states under attended and ignored conditions when compared to passive conditions, and that the attentional modulation under attended and ignored conditions shares a similar direction. Engaging in the task and selectively paying attention to an auditory or visual cue may differentially modulate the auditory cortical neurons. Task engagement may contribute to the same directional components in the attentional modulation of both attended and ignored modalities, whereas attending to one sensory modality may independently modulate the attended and ignored modalities.

## Methods

### Animals

Animal procedures were approved by the Stony Brook University Animal Care and Use Committee and carried out in accordance with the National Institutes of Health standards. Experiments were conducted using male C57BL/6 mice (Charles River Laboratories). Mice were housed with free access to food, but water was restricted after the initiation of behavioral training. On training days, water was available during task performance (2.5 μL for each correct trial); on non-training days, water bottles were provided to the mice for at least 1 hour per day.

### Behavior

Experiments were conducted in a dark, single-walled, sound-attenuating training chamber. The chamber contained three nose pokes, each of which consisted of an infrared LED/infrared phototransistor pair connected to the Bpod system (Sanworks, LLC) for response detection. The activation of a central nose poke was required for trial initiation. One speaker embedded in the wall delivered auditory cues or distractors. Two white LEDs were mounted in two reward nose pokes for the “visual task.” Water rewards were controlled by the Bpod system and delivered from the wall-mounted nose pokes.

Freely moving mice were trained to perform a set of 2AFC tasks, as previously described (Znamenskiy and Zador 2013, Xiong, Znamenskiy et al. 2015). Each trial was initiated when the mouse inserted its nose into the center port of a three-port operant chamber. After a delay period (200–300 ms; uniform distribution), a 100-ms stimulus would be present, indicating which nose poke (left or right) would be rewarded with water. Mice then selected the left or right goal port based on sensory stimuli. Every mouse was trained to perform in a sound block and light block in randomized sequences.

Auditory stimuli consisted of a pseudorandom, 100-ms stream of 30-ms pure overlapping tones presented at 200 Hz. Eighteen possible tone frequencies were logarithmically spaced from 5 to 40 kHz. For each trial, either the low stimulus (5 to 10 kHz) or the high stimulus (20 to 40 kHz) was selected as the target, and the mice were trained to report low or high by choosing the correct port for the water reward. Correct responses were rewarded with water (2.5 μL for each correct trial), and error trials were punished with a 4-s time out. The sound intensity was calibrated to 60 dB.

### Calcium Imaging Procedure

AAV9-calmodulin protein kinase II (CaMKII)-GCaMP6f (University of Pennsylvania Vector Core) was injected into the auditory cortex at the following stereotaxic coordinates: 2.92 mm caudally from bregma, 4.2 mm laterally from midline, and at a 2.25-mm depth from the depth of the bregma. One week after the injection, a prism probe (diameter: 1.0 mm; length: approximately 4.3 mm; pitch: 0.5; numerical aperture: 0.5; Inscopix) was implanted 0.2 mm laterally from the injection site. Three weeks later, a base plate was implanted after checking the calcium signal.

Images were acquired at 20 frames per second using Inscopix nVista. At the beginning of each imaging session, the protective cap was removed from the previously implanted base plate and attached to the microscope. The imaging field of view (maximal size, 1 × 1 mm) was then selected by adjusting the focus. Focal planes were 150–200 μm away from the prism. During recording, the LED output power of the microscope was set at 30% of the maximum. The time stamps of the behavior events were exported from the Bpod system to the microscope for synchronization.

All task and passive sessions were recorded on the same day to prevent a population shift across days. Recording began 15 min after the start of the session, which allowed the mice to switch strategies from those of previous tasks. Each task session was recorded for 10–15 min. The mice had a 1-hour gap with free access to food between the recording sessions to recover both the calcium signal and the motivation of the animals. After the task sessions, the mice had a 1-hour gap with free access to water and food to let them lose motivation before recording the passive sound response. The passive sound response was recorded for 5 min in the same chamber as the task session. The 100-ms cloud of tones was presented every 3–4 s during the passive session.

The acquired images were spatially downsampled by a factor of 2 and corrected for motion using Mosaic (version 1.2; Inscopix, Palo Alto, CA). The spatial and temporal components (Z-scored ΔF/F) of the recorded neurons were extracted from the images using an extended constrained non-negative matrix factorization (CNMF-E) algorithm (Pnevmatikakis, Soudry et al. 2016, Zhou, Resendez et al. 2018); the minimal correlation was set to 0.95 and the minimal peak noise ratio was set to 10 during the initialization step of the CNMF-E. Cell registration was applied to the spatial component, which allowed tracking of the same neuron from different sessions based on the spatial correlation and center distance (Sheintuch, Rubin et al. 2017).

### Data Analysis and Statistics

Criteria for sound responses: If a neuron had at least two consecutive frames within 500 ms after sound onset with *P* values smaller than 0.01 in the bootstrap analysis of a certain trial type, the neuron was identified as a sound response neuron. The baseline was selected between 200–500 ms before sound onset in the passive block, and between 200–500 ms before trial initiation in the task block:

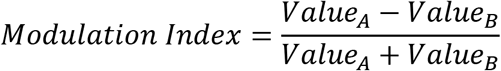

The value is the peak intensity of the average calcium trace within 0–500 ms after sound onset. When the task state is compared with the passive state, A represents the task state, and B represents the passive state. When the attended state is compared with the ignored state, A represents the ignored state, and B represents the attended state.

An SVM was used with a linear kernel for all decoders. The same number of trials (49.00 ± 2.61) from different blocks were randomly selected to balance the decoder. Two-thirds of the data were used for training/validation and the remaining one-third of the data were used for testing. The model was regularized with an L1 penalty to prevent overfitting where the regularization parameter was selected by 5-fold cross-validation. Significant decoding accuracies were determined by comparing the accuracy of the real data with the shuffled data in which behavioral data were shuffled relative to each neuronal activity. All quantified decoding was performed in the full dimensional space. To visualize the high dimensional ensemble activity, we performed principal component analysis on population responses across different attentional conditions (20 trials were selected randomly for each state), then single trial data were projected onto the first three principal components.

The modulation vector from the passive to attended states (***v_PA_***) is defined as the normal vector of the SVM decision boundary at that point from the passive to the attended state, and the modulation vector from the passive to ignored states (***v_PI_***) is defined as the normal vector of the decision boundary at that point from the passive to the ignored state. The angle *θ* between the two modulation vectors is calculated using the following formula:

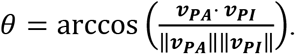

The chance level of *θ* was calculated from the decoder trained using shuffled data.

## Acknowledgements

We thank Drs. Memming Il Park, Anthony Zador, and Yang Yang, and members in Ge’s and Xiong’s laboratories for their critical comments on the manuscript. We thank Dr. Joshua Sanders for assistance with the hardware and software used in the study. We thank Drs. Yang and Ling at the Biostatistical Consulting Core for suggestions concerning data analysis. This work was supported by departmental internal funding and NIH R21DC016746 and R01DC017470 to Q.X.

## Author contributions

P.Y., S.G., and Q.X. designed the experiments. P.Y. performed the experiments. J.S. and L.C. provided technical support. P.Y. and Q.X. analyzed the data and wrote the manuscript.

---

**Supplementary Figure 1.**
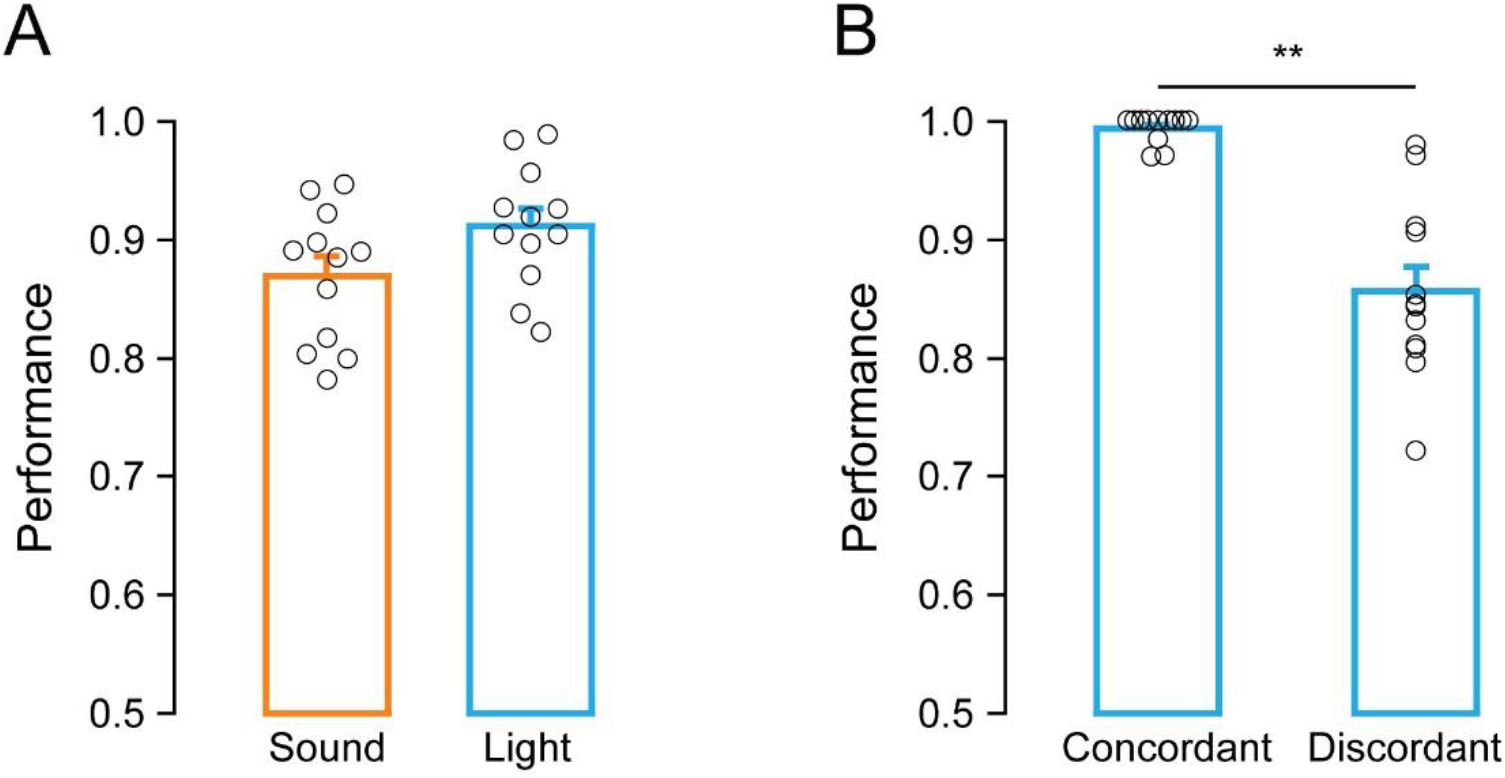
Mice performance in sound block and light block. **A.** Correct response rates from all completed trials in sound blocks and light blocks (n = 5 mice; 12 sessions for each block). **B.** Correct response rates of auditory-visual concordant trials and auditory-visual discordant trials in light blocks. (n = 5 mice; 12 sessions) *P* = 3.96 × 10^−5^, paired-sample *t* test. Error bars: mean ± SEM.

## References

Arieli, A., A. Sterkin, A. Grinvald and A. Aertsen (1996). “Dynamics of ongoing activity: explanation of the large variability in evoked cortical responses.” Science 273(5283): 1868–1871.

Bao, S., V. T. Chan and M. M. Merzenich (2001). “Cortical remodelling induced by activity of ventral tegmental dopamine neurons.” Nature 412(6842): 79–83.

Briggs, F., G. R. Mangun and W. M. Usrey (2013). “Attention enhances synaptic efficacy and the signal-to-noise ratio in neural circuits.” Nature 499(7459): 476–480.

Buran, B. N., G. von Trapp and D. H. Sanes (2014). “Behaviorally gated reduction of spontaneous discharge can improve detection thresholds in auditory cortex.” J Neurosci 34(11): 4076–4081.

Carcea, I., M. N. Insanally and R. C. Froemke (2017). “Dynamics of auditory cortical activity during behavioural engagement and auditory perception.” Nat Commun 8: 14412.

Cohen, M. R. and J. H. Maunsell (2009). “Attention improves performance primarily by reducing interneuronal correlations.” Nat Neurosci 12(12): 1594–1600.

Fritz, J., S. Shamma, M. Elhilali and D. Klein (2003). “Rapid task-related plasticity of spectrotemporal receptive fields in primary auditory cortex.” Nat Neurosci 6(11): 1216–1223.

Hubel, D. H., C. O. Henson, A. Rupert and R. Galambos (1959). “Attention units in the auditory cortex.” Science 129(3358): 1279–1280.

Kastner, S., M. A. Pinsk, P. De Weerd, R. Desimone and L. G. Ungerleider (1999). “Increased activity in human visual cortex during directed attention in the absence of visual stimulation.” Neuron 22(4): 751–761.

Kilgard, M. P. and M. M. Merzenich (1998). “Cortical map reorganization enabled by nucleus basalis activity.” Science 279(5357): 1714–1718.

Kuchibhotla, K. V., J. V. Gill, G. W. Lindsay, E. S. Papadoyannis, R. E. Field, T. A. Sten, K. D. Miller and R. C. Froemke (2017). “Parallel processing by cortical inhibition enables context-dependent behavior.” Nat Neurosci 20(1): 62–71.

Lin, S. C., R. E. Brown, M. G. Hussain Shuler, C. C. Petersen and A. Kepecs (2015). “Optogenetic Dissection of the Basal Forebrain Neuromodulatory Control of Cortical Activation, Plasticity, and Cognition.” J Neurosci 35(41): 13896–13903.

Marlin, B. J., M. Mitre, A. D’Amour J, M. V. Chao and R. C. Froemke (2015). “Oxytocin enables maternal behaviour by balancing cortical inhibition.” Nature 520(7548): 499–504.

Martins, A. R. and R. C. Froemke (2015). “Coordinated forms of noradrenergic plasticity in the locus coeruleus and primary auditory cortex.” Nat Neurosci 18(10): 1483–1492.

Mozolic, J. L., D. Joyner, C. E. Hugenschmidt, A. M. Peiffer, R. A. Kraft, J. A. Maldjian and P. J. Laurienti (2008). “Cross-modal deactivations during modality-specific selective attention.” BMC Neurol 8: 35.

Nelson, A. and R. Mooney (2016). “The Basal Forebrain and Motor Cortex Provide Convergent yet Distinct Movement-Related Inputs to the Auditory Cortex.” Neuron 90(3): 635–648.

Otazu, G. H., L. H. Tai, Y. Yang and A. M. Zador (2009). “Engaging in an auditory task suppresses responses in auditory cortex.” Nat Neurosci 12(5): 646–654.

Petersen, S. E. and M. I. Posner (2012). “The attention system of the human brain: 20 years after.” Annu Rev Neurosci 35: 73–89.

Pnevmatikakis, E. A., D. Soudry, Y. Gao, T. A. Machado, J. Merel, D. Pfau, T. Reardon, Y. Mu, C. Lacefield, W. Yang, M. Ahrens, R. Bruno, T. M. Jessell, D. S. Peterka, R. Yuste and L. Paninski (2016). “Simultaneous Denoising, Deconvolution, and Demixing of Calcium Imaging Data.” Neuron 89(2): 285–299.

Rodgers, C. C. and M. R. DeWeese (2014). “Neural correlates of task switching in prefrontal cortex and primary auditory cortex in a novel stimulus selection task for rodents.” Neuron 82(5): 1157–1170.

Sheintuch, L., A. Rubin, N. Brande-Eilat, N. Geva, N. Sadeh, O. Pinchasof and Y. Ziv (2017). “Tracking the Same Neurons across Multiple Days in Ca 2+ Imaging Data.” Cell Reports 21(4): 1102–1115.

Song, Y. H., J. H. Kim, H. W. Jeong, I. Choi, D. Jeong, K. Kim and S. H. Lee (2017). “A Neural Circuit for Auditory Dominance over Visual Perception.” Neuron.

Vazey, E. M., D. E. Moorman and G. Aston-Jones (2018). “Phasic locus coeruleus activity regulates cortical encoding of salience information.” Proc Natl Acad Sci U S A.

Wimmer, R. D., L. I. Schmitt, T. J. Davidson, M. Nakajima, K. Deisseroth and M. M. Halassa (2015). “Thalamic control of sensory selection in divided attention.” Nature 526(7575): 705–709.

Womelsdorf, T., P. Fries, P. P. Mitra and R. Desimone (2006). “Gamma-band synchronization in visual cortex predicts speed of change detection.” Nature 439(7077): 733–736.

Xiong, Q., P. Znamenskiy and A. M. Zador (2015). “Selective corticostriatal plasticity during acquisition of an auditory discrimination task.” Nature 521(7552): 348–351.

Zhou, P., S. L. Resendez, J. Rodriguez-Romaguera, J. C. Jimenez, S. Q. Neufeld, A. Giovannucci, J. Friedrich, E. A. Pnevmatikakis, G. D. Stuber, R. Hen, M. A. Kheirbek, B. L. Sabatini, R. E. Kass and L. Paninski (2018). “Efficient and accurate extraction of in vivo calcium signals from microendoscopic video data.” Elife 7.

Zimmer, U., S. Itthipanyanan, T. Grent-’t-Jong and M. G. Woldorff (2010). “The electrophysiological time course of the interaction of stimulus conflict and the multisensory spread of attention.” Eur J Neurosci 31(10): 1744–1754.

Znamenskiy, P. and A. M. Zador (2013). “Corticostriatal neurons in auditory cortex drive decisions during auditory discrimination.” Nature 497(7450): 482–485.

